# Structural Analyses of a GABARAP∼ATG3 Conjugate Uncover a Novel Non-covalent Ubl-E2 Backside Interaction

**DOI:** 10.1101/2024.08.14.607425

**Authors:** Kazuto Ohashi, Takanori Otomo

**Affiliations:** Department of Integrative Structural and Computational Biology, The Scripps Research Institute, 10550 North Torrey Pines Rd, La Jolla, CA 92037, USA; Institute for Molecular and Cellular Regulation, Gunma University, 371-8512 Gunma, Japan; San Diego Biomedical Research Institute, 3525 John Hopkins Ct, San Diego, CA 92121, USA

**Author notes:** Correspondence: Takanori Otomo.

## Abstract

Members of the ATG8 family of ubiquitin-like proteins (Ubls) are conjugated to phosphatidylethanolamine (PE) in the autophagosomal membrane, where they recruit degradation substrates and facilitate membrane biogenesis. Despite this well-characterized function, the mechanisms underlying the lipidation process, including the action of the E2 enzyme ATG3, remain incompletely understood. Here, we report the crystal structure of human ATG3 conjugated to the mammalian ATG8 protein GABARAP via an isopeptide bond, mimicking the Ubl∼E2 thioester intermediate. In this structure, the GABARAP∼ATG3 conjugate adopts an open configuration with minimal contacts between the two proteins. Notably, the crystal lattice reveals non-covalent contacts between GABARAP and the backside of ATG3’s E2 catalytic center, resulting in the formation of a helical filament of the GABARAP∼ATG3 conjugate. While similar filament formations have been observed with canonical Ub∼E2 conjugates, the E2 backside-binding interface of GABARAP is distinct from those of Ub/Ubl proteins and overlaps with the binding site for LC3 interacting region (LIR) peptides. NMR analysis confirms the presence of this non-covalent interaction in solution, and mutagenesis experiments demonstrate the involvement of the E2 backside in PE conjugation. These findings highlight the critical role of the E2 backside in the lipidation process and suggest evolutionary adaptations in the unique E2 enzyme ATG3.

## INTRODUCTION

Autophagy is an intracellular process in which the double-membrane autophagosome transports cytoplasmic materials to lysosomes for degradation and recycling^1–4^. While starvation-induced autophagy is generally non-selective, autophagy can also selectively target damaged materials or large cytotoxins, such as aberrant organelles and invading bacteria. The ATG8 family ubiquitin-like proteins (Ubls) play crucial roles in autophagy by localizing to the autophagosomal membrane and recruiting specific substrates through interactions with a tetra-amino acid motif, known as ATG8-interacting motif (AIM), LC3-interacting regions (LIR), or LC3 recognition sequence (LRS) of “autophagy receptor” proteins^5–9^. In yeast, a single gene, Atg8, is responsible for substrate recruitment, whereas mammalian cells utilize six ATG8 paralogues (LC3A, LC3B, LC3C, GABARAP, GABARAPL1, and GABARAPL2) to collectively cover a wide range of substrates. Additionally, Atg8 can perturb membrane morphology by inserting its aromatic residues into the membrane, thereby facilitating the expansion of autophagosomal membranes in yeast^10–14^. Similarly, LC3 and GABARAP proteins are also essential for the efficient formation of autophagosomes in mammalian cells^10, 15, 16^.

These functions of ATG8 proteins depend on their covalent attachment to phosphatidylethanolamine (PE) molecules in the autophagosomal membrane^17^. After protein synthesis, ATG8 proteins are cleaved by the ATG4 protease, exposing a glycine residue as the new C-terminus^18^. The processed ATG8 is activated by the E1 enzyme ATG7 and transferred to the E2 enzyme ATG3, forming the ATG8∼ATG3 thioester intermediate. While these initial reactions occur constitutively within cells, the subsequent transfer of ATG8 from ATG3 to PE is specifically stimulated upon the induction of autophagy, a process that requires the involvement of the ATG12-ATG5-ATG16 E3 complex^19, 20^.

Previous structural studies on yeast and plant Atg7 proteins revealed the molecular mechanisms of the first two steps of the lipidation, specifically the activation of ATG8 by ATG7 and the subsequent transfer of ATG8 from ATG7 to ATG3^21–26^. By contrast, the mechanism of the PE conjugation step is still elusive due to the lack of structural data representing intermediate states. Additionally, the unique molecular features of ATG3 and the ATG12-ATG5-ATG16 E3 complex complicate the inference of their mechanisms based on knowledge of canonical Ub/Ubl E2 and E3 enzymes^27^.

ATG3 belongs to one of the 17 classes of the Ub/Ubl E2 enzyme superfamily^28^, and its architecture shows moderate differences from canonical E2 enzymes^29^. While canonical E2s typically consist of ∼150 residues with four α-helices and four β-strands, the catalytic domain of ATG3 is larger, with ∼185 residues, and includes two additional α-helices and two β-strands. Furthermore, the residues in the catalytic center of ATG3 differ from those of canonical E2s. For example, the asparagine residue that stabilizes the thioester linkage^30^ and the aspartate and tyrosine residues that de-solvate the substrate amino group^31^ in the canonical E2s are not conserved in ATG3^29^. It has been proposed that threonine 213 in yeast Atg3 may perform the role of the asparagine residue in canonical E2s^32^, but corresponding residues for the aspartate and tyrosine in ATG3 have not been identified. The sequence following the catalytic cysteine residue in ATG3 forms a unique element known as the handle region (HR), which consists of Helix F–one of the extra α-helices–and a short loop^29^. The HR of yeast Atg3 contains an AIM, which plays a role in its membrane localization^33^, while this motif is absent in mammalian ATG3.

In addition to differences in the catalytic domain, full-length ATG3 contains two disordered regions: the N-terminal ∼25 residues and the ∼100 residue-long insertion between β-strands 2 and 3, referred to as the flexible region (FR) ^29, 34^. The N-terminus serves as the membrane anchor domain by folding into an amphipathic helix on membranes^35–39^. The FR plays crucial roles in E1 and E3 interactions^24, 25, 29, 40, 41^. In human ATG3, residues 157-181 near the C-terminal end of the FR bind to ATG7^42^, while residues 140-170 bind to ATG12 in the E3 complex^43^. The overlap between the E1- and E3-binding regions suggests that the E1-E2 and E2-E3 interactions are mutually exclusive^40^, a common feature among the Ub/Ubl cascades^44^. Similarly, in the yeast *Saccharomyces cerevisiae* (Sc) Atg3, Helix C in the FR is responsible for both of E1- and E3-binding^21, 24, 25, 29, 41, 45^. Moreover, ScAtg3’s Helix C interacts with its own catalytic domain to suppress its activity^29^, earning it the designation of the E1, E2, E3-interacting region (E123IR)^45^. However, whether the FR of mammalian ATG3 also possess a suppressive activity remains unknown. The FR also can interact with Atg8 and LC3/GABARAP proteins, participating in efficient conjugation reaction as well as membrane reorganization^41, 46^.

The catalytic domain of ATG3 has been visualized in two distinct conformations. In the apo form of the full-length ScAtg3, the E123IR is bound to a site composed of Helices G and F, which is distant from the catalytic core^29^. This interaction allosterically stabilizes an inactive “closed” conformation, where the catalytic cysteine is buried^45^. Sequestration of E123IR from the catalytic domain, either by E1 or E3 binding or by deletion of E123IR, induces a conformational change that extends Helix F and exposes the catalytic cysteine^24, 25, 45^. Although this E123IR-unbound “open” conformation of the catalytic domain is likely closer to an active form, it has been suggested that additional conformational changes may be required for ATG8– PE conjugation activity^32, 45^.

To gain deeper insights into the PE conjugation process, we performed structural analyses of an isopeptide-stabilized human GABARAP∼ATG3 intermediate mimic (∼ denotes a thioester linkage or its mimic). We determined the crystal structure of a GABARAP∼ATG3 conjugate, which appears to represent an inactive conformation. Unexpectedly, the crystal packing revealed a previously unobserved non-covalent interaction between GABARAP and ATG3. Point mutations at this contact surface on ATG3–located on the opposite side of the catalytic core, referred to as the “backside”–disrupted the non-covalent GABARAP interaction and impaired PE conjugation. These findings underscore the critical role of the ATG3 backside in PE lipidation process.

## RESULTS AND DISCUSSION

### Crystal Structure of a GABARAP∼ATG3 Intermediate Mimic

The thioester bond between ATG8 and ATG3 is labile^19^, which has hindered structural studies of the ATG8∼ATG3 intermediate. To stabilize this linkage, we replaced the thioester with an isopeptide bond by conjugating GABARAP to a human ATG3 mutant with the catalytic cysteine residue (Cys264) mutated to lysine. This strategy has been used previously to stabilize Ub∼E2 intermediates in structural studies^47^. Although Lys264 is not the natural substrate of the E1 enzyme ATG7, human ATG7 was able to attach GABARAP to ATG3^C264K^ (Fig. S1).

We attempted to crystallize the resulting GABARAP∼ATG3^C264K^ conjugate but were unsuccessful. To facilitate crystallization, we deleted three flexible regions of ATG3: the N-terminal 26 residues, and the first 56 and the last 11 residues of the FR, none of which are involved in either E1 or E3 interaction^29, 42, 43^. This shorter construct, referred to as ATG3_crystal_ (Fig. 1A), was able to conjugate with GABARAP (Fig. S1). The resultant GABARAP∼ATG3_crystal_ conjugate crystallized successfully. The crystals diffracted synchrotron X-rays to 2.7 Å, and the structure was determined to the R and Rfree factors of 0.219 and 0.248, respectively (Table S1). Most residues, except those of the FR, are visible in the electron density map. The electron density of the lysine side chain at position 264 in the catalytic site of ATG3 is connected, albeit weakly, to the C-terminal Gly116 of GABARAP (Fig. S2). This confirms the conjugation between the two proteins.

**Figure 1.**
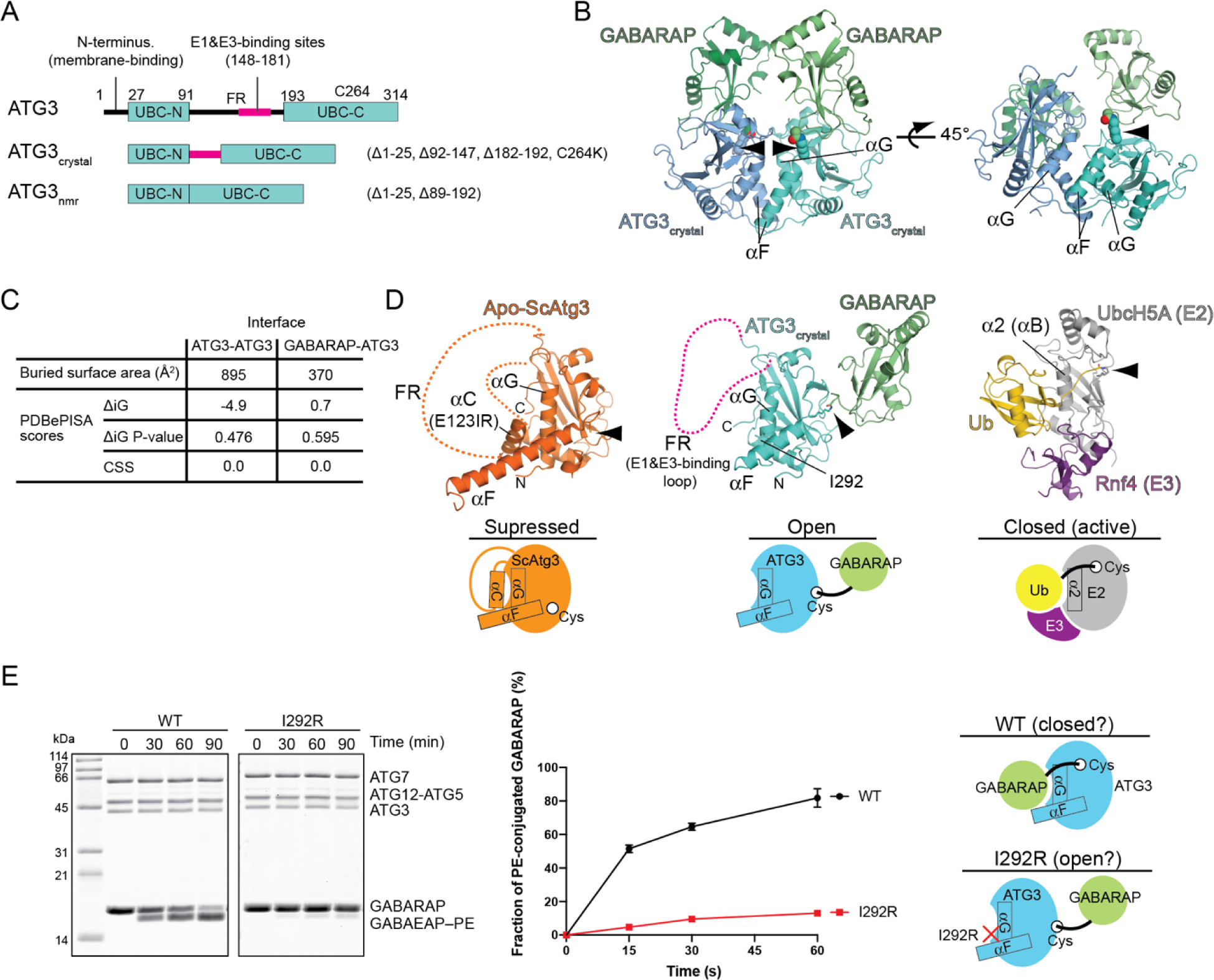
Crystal structure of the GABARAP∼ATG3 conjugate mimic. (A) The primary structure of human ATG3 and the constructs used in this study. (B) The crystal structure of the GABARAP∼ATG3_crystal_ isopeptide conjugate. The two molecules of the GABARAP∼ATG3_crystal_ conjugate in the asymmetric unit are shown. The covalent linkages between Lys264^ATG3^ and Gly116^GABARAP^ are indicated by arrowheads. (C) Metrics from a PDBePISA analysis of the molecular interfaces within the asymmetric unit. (D) Structural comparisons of the GABARAP∼ATG3_crystal_ conjugate (middle) with apo ScAtg3 (left, PDB ID: 2DYT) ^29^ and the Ub∼Ubc5HA (E2)-Rnf4 (E3) complex (right, PDB ID: 1Z5S])^53^. Cartoons depict the state of each structure. (E) GABARAP–PE conjugation assay with wild-type and I292R mutant ATG3. SDS-PAGE analysis of the reaction is shown on the left. ATG16N is not visible on the SDS-PAGE due to its small size. The quantification of the results is shown on the right. Data points are presented as the mean ± S.D. of three independent experiments. Putative configurational states of the GABARAP∼ATG3 conjugate are depicted as cartoons.

The asymmetric unit contains two of GABARAP∼ATG3_crystal_ molecules, related by two-fold non-crystallographic symmetry (NCS), which pack against each other through contacts between the two ATG3 molecules (Fig. 1B). PDBePISA interface analysis reported a complex formation significance score (CSS) of 0.0, suggesting that this interface, which buries a surface area of 895 Å^2^, is crystallographic (Fig. 1C). The interface consists of Helices F and G in the C-terminal region and overlaps with the E123IR (Helix C)-binding surface of ScAtg3 (Fig. 1D)^29^. The conjugated GABARAP is positioned in front of the catalytic site of ATG3, burying a small surface area of ∼370 Å^2^ between the two proteins. PDBePISA analysis also suggests that this interface, mediated mostly by the C-terminal residues of GABARAP, is energetically unfavorable, with a CSS of 0.0 (Fig. 1C and 1D). The isopeptide linkage is exposed to solvent and has no contacts with other residues in the catalytic center (Fig. 1D). Thus, it appears that the relative position between GABARAP and ATG3 is dictated by crystal packing, similar to the previous crystallization of the Ub∼Ubc5Hb intermediate^48, 49^.

In solution, canonical Ub/Ubls attached to E2 enzymes are flexible, which prevents substrate proteins from attacking the thioester linkage^48, 50, 51^. In contrast, in their E3-activated forms, the C-terminal residues of Ub/Ubls and the thioester linkages make extensive contacts with the residues in the E2 catalytic centers^51–56^; the Ub folds are fixed on the E2 surface, composed of the second helix (termed α2 or αB) and its surrounding residues, generating the “closed” Ub/Ubl∼E2 configuration (Fig. 1D). Thus, the GABARAP∼ATG3_crystal_ structure likely represents an inactive “open” configuration.

Since the FR was not visible in the crystal and the putative FR-binding surface of the catalytic domain was blocked by another ATG3 molecule, we considered whether the FR was displaced from the catalytic domain upon crystallization. Such displacement might occur if the FR associates weakly with the catalytic domain, as is the case with ScAtg3, where the E123IR and the catalytic domain of ScAtg3 associate in trans with a Kd of ∼200 µM^45^. Alternatively, the catalytic domain-binding region of the FR in human ATG3 might be within the region that was deleted to facilitate crystallization. To explore these possibilities, we performed NMR experiments using ^15^N-labeled ATG3^FR^. As reported previously, the ^1^H-^15^N HSQC spectrum of ATG3^FR^ shows a pattern of disordered fragments^43^. When a 2.5-fold excess of the unlabeled ATG3^ΔFR^ was added, the spectrum remained unchanged (Fig. S3), suggesting that no region of the FR in human ATG3 binds to the catalytic domain with detectable affinity. Therefore, when the FR is not engaged with E1 or E3, it is likely disordered, which explains the lack of the electron density for the FR. This finding suggests that the regulatory mechanism of the enzyme might differ between ScAtg3 and human ATG3.

Since the “closed” configuration of the GABARAP∼ATG3 conjugate was not observed in the crystal, we considered whether the conjugate might adopt such a configuration during the reaction. As previously suggested for the yeast system, ATG8 could be bound to the surface composed of Helices F and G^45^. To test this configuration, we introduced a mutation on Helix G (I292R) designed to disrupt the expected docking of GABARAP. The I292R mutant ATG3 failed to produce GABARAP–PE conjugate (Fig. 1E), despite being properly loaded with GABARAP by ATG7 (Fig. S4). Thus, the GABARAP–PE conjugation requires the surface including Helix G, supporting the existence of the active “closed” configuration.

### The ATG3 Catalytic Center Adopts an Open Conformation

The catalytic center of GABARAP-loaded ATG3_crystal_ adopts an “open” conformation that is overall very similar to those observed in Arabidopsis thaliana (At)Atg3 bound to Atg7 and in ScAtg3 conjugated to ScAtg7^24, 25^ (Fig. 2A–2D). For instance, the GABARAP-conjugated catalytic residue (C264K) and Thr244, which corresponds to Thr213 in ScAtg3 and is suggested to stabilize the ScAtg8-ScAtg3 thioester linkage^32^, are superimposable with corresponding residues in these previous structures. Unlike in the activated form of canonical E2 enzymes, C264K is exposed and located far (∼9 Å) from Thr244 (Fig. 2A), suggesting that ATG3 in the crystal is not in the active state.

**Figure 2.**
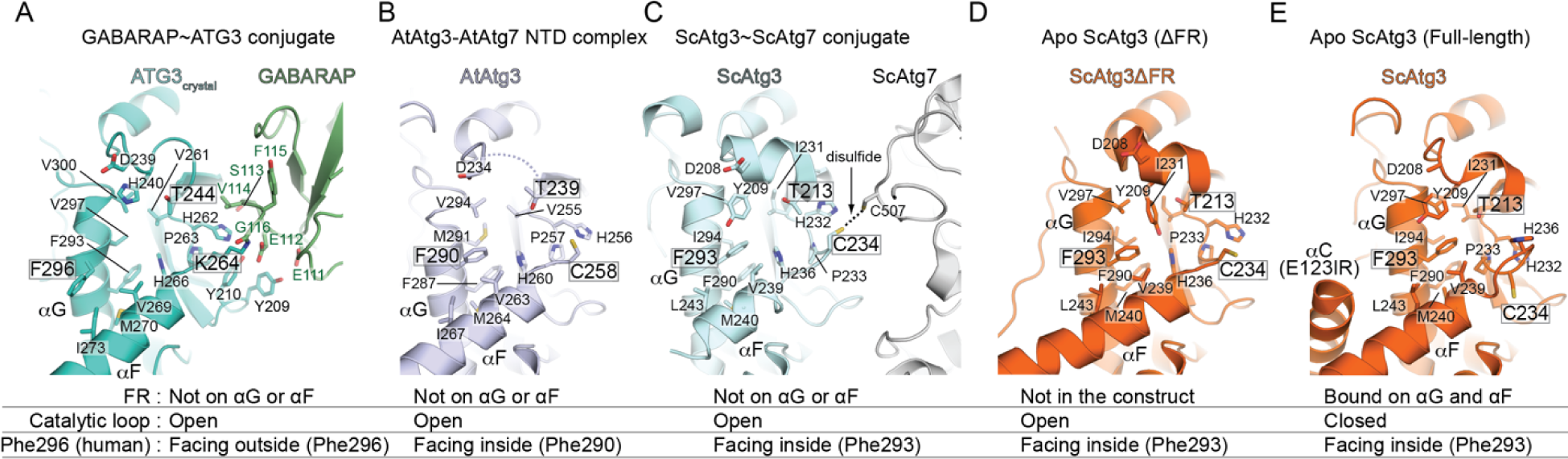
The catalytic centers of ATG3/Atg3 structures. (A) The GABARAP∼ATG3 isopeptide conjugate. (B) The AtAtg3-AtAtg7 NTD complex (PDB ID: 3VX8)^25^. (C) ScAtg3∼ScAtg7 disulfide-crosslinked conjugate (PDB ID: 4GSL)^24^. (D) Apo ScAtg3ΔFR (PDB ID:6OJJ)^45^. (E) Apo ScAtg3 full-length (PDB ID: 2DYT) ^29^. The states of the FR, catalytic loop, and Phe296 (Phe290/Phe293) are described at the bottom. The residues mentioned in the text are labeled in boxes.

A unique feature of the GABARAP-loaded ATG3_crystal_ is the side chain rotamer conformation of Phe296 on Helix G (Fig. 2A). In the previous structures, the corresponding residues (Phe290 in AtAtg3 and Phe293 in ScAtg3) face inward and make contacts with residues in the protein core, such as Phe287/290, His260/236, Val263/239 and Val294/297 (the residue numberings for AtAtg3/ScAtg3) (Fig. 2B–2E). However, in the GABARAP-loaded ATG3_crystal_, the phenyl ring of Phe296 flips out of this hydrophobic core and makes new contacts with lle273 of Helix F on the protein surface (Fig. 2A). Phe296 and Ile273 also make contacts with the corresponding residues of the NCS copy, which might suggest that the conformation of Phe296 is a consequence of crystallization.

Phe293 of ScAtg3 has been suggested to act as a suppressor of PE-conjugation activity; the F293S mutation of ScAtg3 facilitated ScAtg8 transfer to PE even in the absence of the E3^32^. Because the loss of the phenyl ring from the originally observed inward-facing position is a shared feature between the F293S mutation of ScAtg3 and our structure, we wondered if the flipping of the phenyl ring induces catalysis. However, it has been recently reported that the same mutation in human ATG3 (ATG3^F296S^) impairs PE conjugation activity^57^, and we also confirmed this observation (Fig. S5). Since ATG7-mediated GABARAP loading onto ATG3^F296S^ was as efficient as it was onto wild-type ATG3 (Fig. S4), ATG3^F296S^ appears to be incapable of transferring GABARAP to PE. This may be because the phenyl ring plays a positive role in stabilizing the active Ubl∼E2 configuration in the human system. These findings suggest a distinct mechanism in human ATG3 that warrants further investigation to fully understand its role in PE conjugation.

### GABARAP∼ATG3 Packs into a Filament in the Crystal

In the crystal, GABARAP makes contacts with an ATG3 molecule conjugated to a different GABARAP. The repetition of this inter-conjugate association leads to the formation of a filament composed of the GABARAP∼ATG3 conjugate (Fig. 3A). Such filamentous crystallization has been observed with several canonical Ub∼E2 conjugates^48, 49^. Similar to the Ub∼Ubc5Hb filament, the GABARAP∼ATG3 filament is helical, although the former filament is right-handed, whereas the latter is left-handed (Figs. 3A and 3B)

**Figure 3.**
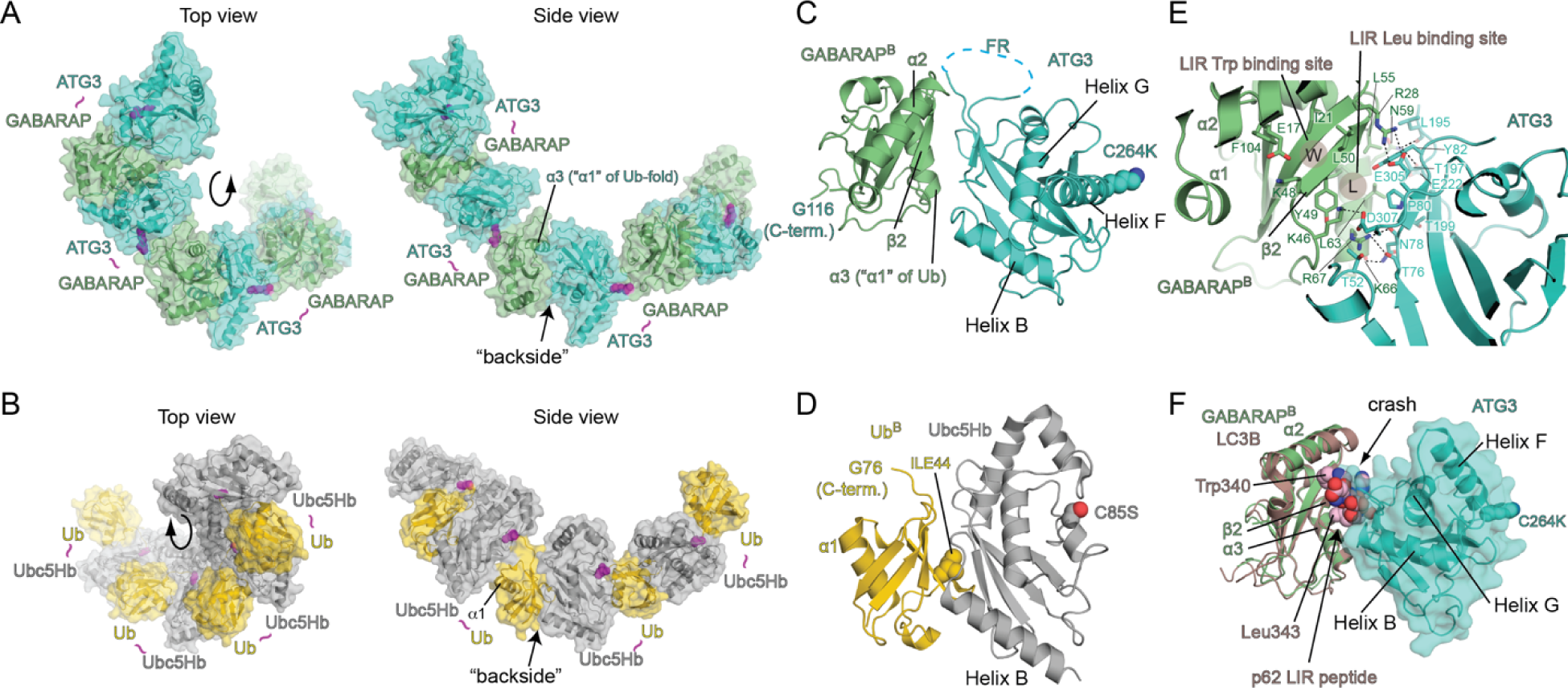
Filamentous assembly of the GABARAP∼ATG3 conjugate in the crystal. (A, B) Comparison of the GABARAP∼ATG3_crystal_ (A) and Ub∼Ubc5Hb (PDB ID: 3A33) (B) filaments in crystals. (C, D) Structures of the non-covalent ATG3-GABARAP^B^ (C) and Ubc5Hb-Ub^B^ (D) pairs shown in an orientation where ATG3 and Ubc5Hb are structurally aligned. (E) Close-up view of the non-covalent interface between ATG3 and GABARAP^B^. (F) Superimposition of the LC3B-p62 LIR peptide complex (PDB ID: 2ZJD) on the structure of the non-covalent ATG3-GABARAP^B^ pair. The p62 LIR peptide, shown in sphere, sterically crashes into the ATG3 backside.

A structural comparison of the non-conjugated ATG3 and GABARAP with Ubc5Hb and Ub reveals that both GABARAP and Ub bind to the opposite side of their respective E2 enzymes’ catalytic centers, a region known as the “backside” (Fig. 3C and 3D). However, the surfaces of Ub and GABARAP involved in these contacts are different. Ub’s surface is centered around Ile44 (Fig. 3D), a residue commonly involved in the interactions with a broad range of Ub-binding domains^58, 59^. In contrast, GABARAP binds to ATG3 through a surface opposite to the residue corresponding to Ile44 in Ub, which consists of α-helix 3 and the residues flanking β-strand 2 (Fig. 3C).

The space between α-helix 3 and β-strand 2 forms a hydrophobic pocket that accommodates the second hydrophobic residue of the LIR motif (Fig. 3E). While the pocket is not occupied by any residues of ATG3, the surrounding residues contact ATG3, leaving insufficient space to accommodate a LIR peptide (Fig. 3E and 3F). A PDBePISA analysis reports a CSS of 0.161 and ΔiG of −2.6 kcal/mol for the ATG3-GABARAP^B^ (^B^; E2 backside-bound) contact. This suggests that this non-covalent interface is energetically favorable, more so than the covalent interface within the GABARAP∼ATG3 conjugate as described above. The ATG3-GABARAP^B^ contact buries a surface area of ∼700 Å^2^, which is comparable to that between UbcH5 and Ub^B^.

### GABARAP Interacts Weakly with the “Backside” of ATG3

The Ub∼UbcH5 conjugate self-associates not only in crystals but also in solution through the same backside interaction^60^. We wondered if the GABARAP∼ATG3 conjugate might also self-associate in solution. To investigate this possibility, we used NMR spectroscopy, similar to the approach in the previous study of the Ub∼UbcH5c polymerization^60^. We generated the GABARAP∼ATG3_crystal_ conjugate in which only GABARAP was labeled with ^15^N. The ^1^H-^15^N HSQC spectrum of this sample showed almost no signals (Fig. 4A), indicating the self-association of the GABARAP∼ATG3 conjugates in solution.

**Figure 4.**
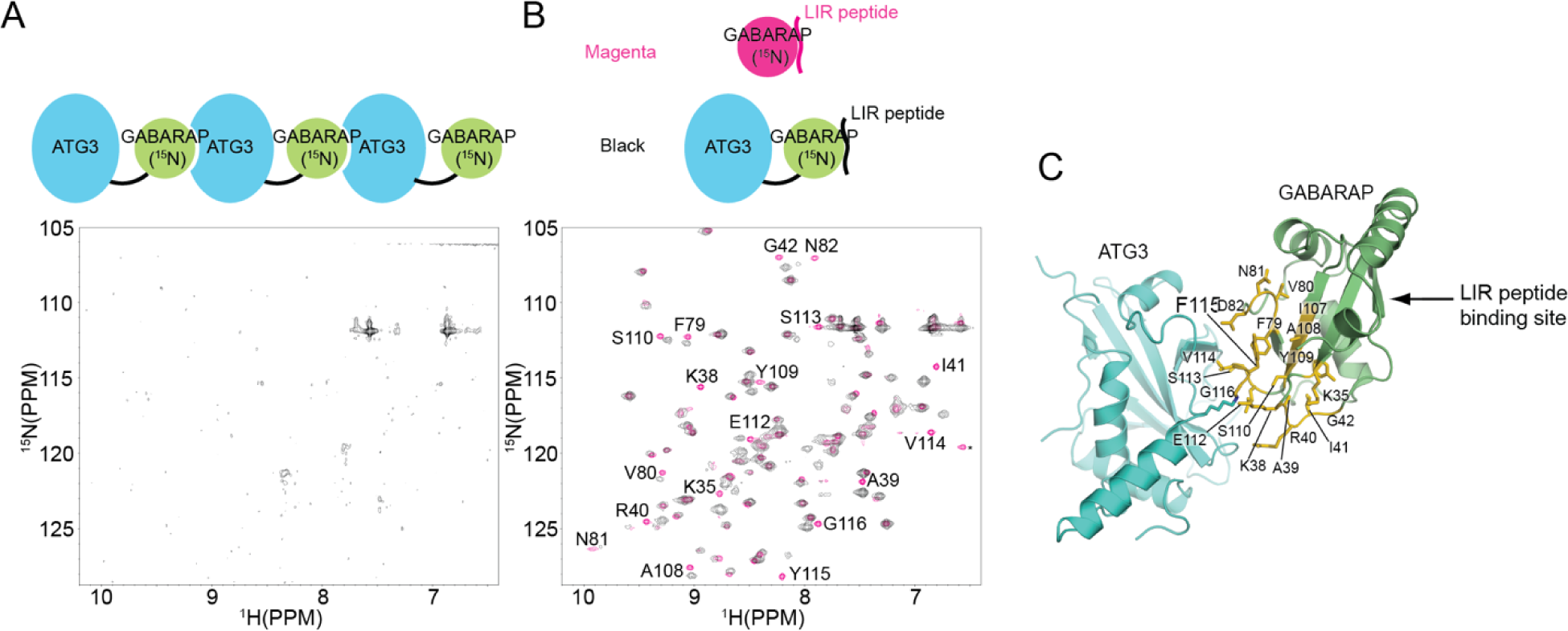
The GABARAP∼ATG3 conjugate interacts in solution. (A, B) ^1^H-^15^N HSQC spectrum of the GABARAP (^15^N-labeled)∼ATG3 (non-labeled) conjugate in the absence (A) or the presence (B) of p62 LIR peptides (non-labeled). Both spectra are shown in black. Also shown in (B) is the spectrum of the non-conjugated ^15^N-GABARAP in complex with p62 LIR peptide (non-labeled) in magenta. The magenta peaks that shift or are absent in the black spectrum are labeled with their assignments. (C) The residues labeled in (B) are shown in yellow and as sticks representation in the GABARAP∼ATG3 crystal structure.

As mentioned earlier, the ATG3-GABARAP^B^ interface observed in our crystal is structurally incompatible with LIR peptide binding (Fig. 3E). Therefore, we hypothesized that the addition of a LIR peptide to the NMR sample could disrupt the ATG3-GABARAP^B^ inter-conjugate association and thereby restore the peaks. As shown in Fig. 4B, the addition of the p62 LIR peptide resulted in the appearance of all GABARAP resonances. Comparison of the LIR peptide-GABARAP∼ATG3 complex spectrum with that of LIR peptide-GABARAP complex shows that the peaks shifting between the two correspond to residues adjacent to the conjugated ATG3 in the crystal structure (Fig. 4C). The peaks of residues in the LIR-binding surface of GABARAP remain unaffected, indicating that the LIR peptide binds to GABARAP of the GABARAP∼ATG3 conjugate. These data suggest that the LIR-binding surface of GABARAP associates non-covalently with ATG3.

Next, we sought to confirm that ATG3 interacts with GABARAP through its backside surface. Because the disordered N-terminus and FR would hamper NMR analyses of ATG3 by generating too many overlapping intense peaks, we created a new construct, ATG3_nmr_, that lacks these disordered regions (Fig. 1A). The ^15^N-labeled ATG3_nmr_ yielded a high-quality ^1^N-^15^N HSQC spectrum, and upon titration of free unlabeled GABARAP into this sample, a number of peaks shifted gradually or broadened (Fig. 5A). Fitting the titration curves yielded similar Kd values, indicating a single binding mode, with an average Kd of 1.3 ± 0.3 mM (Fig. 5B). The affected resonances are concentrated on the ATG3 backside including the side chain of Asn78 (Fig. 5C-5E), which, in the crystal structure, forms a hydrogen bond with the backbone carbonyl group of Lys66 on α-helix 3 of GABARAP (Fig. 3E). These data confirm that ATG3 interacts with GABARAP through its backside in the manner observed in the crystal structure.

**Figure 5.**
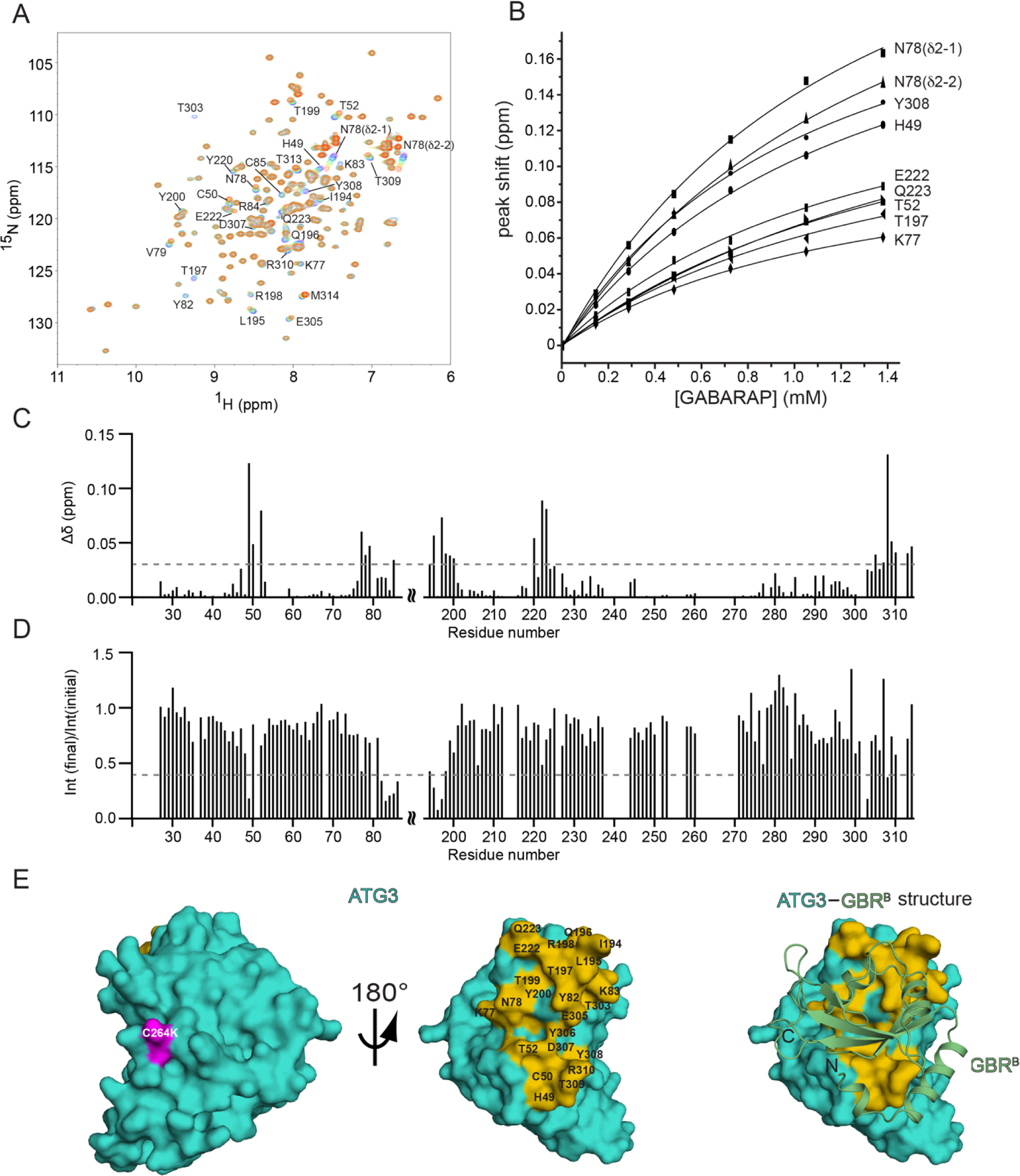
GABARAP interacts non-covalently with the backside of ATG3’s catalytic site in solution. (A) Overlay of HSQC spectra of ^15^N-labeled ATG3_nmr_ titrated with unlabeled GABARAP. The peaks that shifted or broadened upon the addition of GABARAP are labeled with their residue assignments. (B) Titration curves of the chemical shift changes of peaks that shifted more than 0.06 ppm are shown with fitted curves. (C) Plot of the combined ^1^H and ^15^N chemical shift changes of ATG3 peaks as a function of residue number. (D) Plot of ratios of ^1^H-^15^N peak intensities of the final GABARAP titration point compared to the initial ones. (E) Chemical shift perturbation plotted on each residue of ATG3 structure. The residues whose peak shifted more than 0.03 ppm (above the dotted line in (C)) or lost signal intensity by more than 60% (below the dotted line in (D)) are colored yellow and labeled on the surface of ATG3. The catalytic residue C264 is shown in magenta and labeled as its mutated form (C264K). On the right, the non-covalently contacting pair of ATG3 and GBR^B^ in the crystal structure is shown.

### The ATG3 Backside is Critical for Lipidation

To assess the importance of the backside of ATG3 in ATG8 lipidation, we introduced point mutations to disrupt this interaction. Since ATG3 associates with the E1 ATG7 not only through the FR but also via its backside^24, 25^, and this backside binding positions the catalytic cysteine optimally for ATG8 transfer from ATG7 and ATG3, these mutations had to avoid disrupting the ATG3-ATG7 interface. We targeted Thr52 and Thr197, located at the edge of the ATG3-ATG7 interface. Their side chains are surrounded by GABARAP residues in our structure but are much less so by Atg7 in the AtAtg7-AtAtg3 and ScAtg7∼ScAtg3 structures (Fig. S6).

The ^1^N-^15^N HSQC spectra of the T52R and T197R ATG3_nmr_ mutants are very similar to that of the wild-type (Fig. 6A), suggesting that these mutations did not alter the overall protein structure. However, adding unlabeled GABARAP to these samples induced no chemical shift perturbation (Fig. 6A), indicating that the mutations prevent GABARAP binding, at least at the highest concentration achieved in the titration. Both mutations severely diminished GABARAP lipidation (Fig. 6B) without affecting GABARAP loading by ATG7 (Fig. S4), indicating that the backside of ATG3 is involved in the PE attachment step.

**Figure 6.**
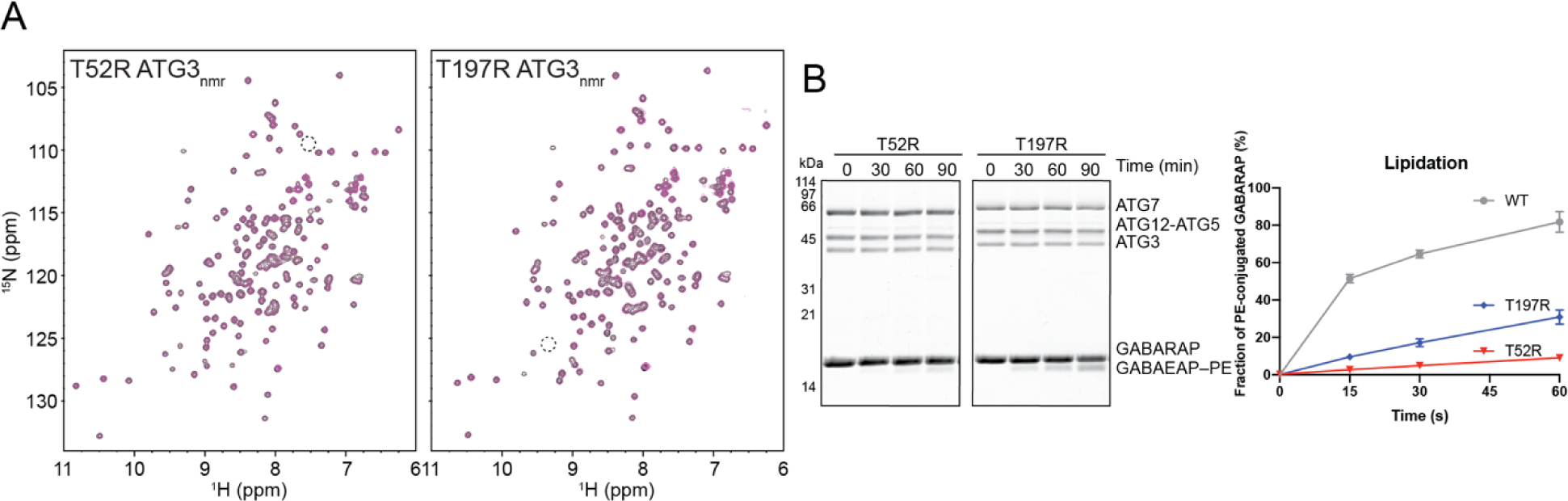
The ATG3 backside is important for GABARAP–PE conjugation. (A) Overlays of ^1^H-^15^N HSQC spectra of ATG3_nmr_^T52R^ (left) and ATG3_nmr_^T197R^ (right) without (black peaks) and with GABARAP (red peaks) are shown. The positions of the Thr52 and Thr197 amide resonances in the wild-type spectrum (Fig. 5A) are indicated by dotted circles. (B) GABARAP–PE conjugation by the mutants. The data and quantification are presented in the same manner as in Fig. 1E.

Notably, Thr52 corresponds to Ser22 in UbcH5c, a residue whose mutation to Arg (S22R) is known to disrupt the non-covalent Ub interaction^60^. Thus, despite differences between the Ub and GABARAP surfaces mediating E2 backside interactions, similar mutations have similar effects, suggesting that the backside interaction of ATG3 may be evolutionarily related to that of canonical E2 enzymes.

### Role of ATG3 Backside Interaction

Previous studies on canonical Ub/Ubl pathways have shown that the self-association of Ub∼E2 intermediates through the E2 backside promotes Ub/Ubl chain formation in a processive manner ^60–63^. While the precise role of the ATG3 backside interaction remains unclear, and GABARAP does not self-conjugate to form such chains, an analogy could be drawn.Once a GABARAP–PE conjugate is produced, it may help recruit another GABARAP∼ATG3 conjugate to the membrane, thereby facilitating the production of GABARAP–PE conjugates on the same membrane.

As mentioned earlier, the ATG3-GABARAP^B^ interaction has a millimolar-range affinity, which is weaker than the interactions between Ub E2s and Ub^B^ (high micromolar range) ^60, 64^ and between Ubc9 and SUMO^B^ (nanomolar range)^61–63, 65, 66^. Thus, recruiting the GABARAP∼ATG3 intermediate solely through the ATG3-GABARAP^B^ interaction may not be efficient. Instead, this interaction could potentially cooperate with other mechanisms, such as recruitment by the ATG12-ATG5-ATG16L1 E3 complex through the ATG3-ATG12 interaction^43^ and ATG3’s intrinsic affinity for highly curved membranes^35–38, 57^. A precise understanding of the ATG3 backside interaction and its role in the lipid conjugation cascade will require further research aimed at dissecting the complex network of interactions involved in the human ATG conjugation system.

## Acknowledgments

We thank Dr. Gerard Kroon for assistance with NMR experiments and Chinatsu Otomo for help with biochemical assays. This work was supported by a grant from the National Institute of General Medical Sciences (GM092740) to T.O. K.O. was a recipient of the Japan Society for the Promotion of Science Overseas Fellowship. Use of the Stanford Synchrotron Radiation Lightsource, SLAC National Accelerator Laboratory, is supported by the U.S. Department of Energy, Office of Science, Office of Basic Energy Sciences under Contract No. DE-AC02-76SF00515. The SSRL Structural Molecular Biology Program is supported by the DOE Office of Biological and Environmental Research, and by the National Institutes of Health, National Institute of General Medical Sciences (P30GM133894).

## Methods

### Protein expression and purification

The DNA fragments coding the human ATG3 variants (full-length, 1-314; ATG3_crystal_, residues 27–91+148–181+193–314 with the C264K mutation; ATG3_nmr_, residues 24–88+192–314) were cloned into a modified pGEX vector containing a TEV protease-cleavable N-terminal glutathione-S-transferase (GST) tag. *E. coli* BL21(DE3) cells transformed with these plasmids were grown in LB media, and protein expression was induced by adding 0.2 mM IPTG when OD_600_ reached 0.8. The cells were harvested after overnight at 22°C. The cells were resuspended in 20 mM Tris pH8.0, 300 mM NaCl, 1 mM DTT, 1 mM EDTA and lyzed using a C3 cell homogenizer (Avestin). The lysate was clarified by centrifugation at 40,000 x g for 30 min and the supernatant was mixed with glutathione sepharose resin (Gold Bio). After a 30 min incubation, the supernatant was discarded and the resin was washed with 20 mM Tris pH 8.0, 500 mM NaCl, 1 mM DTT, 1 mM EDTA, 0.2% TritonX-100 and then re-washed with 20 mM Tris pH 8.0, 300 mM NaCl, 1 mM DTT, 1 mM EDTA. The proteins were eluted in 20 mM Tris pH 8.0, 300 mM NaCl, 1 mM DTT, 1 mM EDTA, and 15 mM glutathione. The eluted GST-ATG3 was digested with TEV protease at room temperature. The digested proteins were further purified by Source 15Q anion exchange chromatography and size exclusion chromatography. The purified proteins were concentrated and frozen for storage until use.

The DNA sequence coding human GABARAP was cloned into a modified pET (Novagen) vector with a TEV-cleavable N-terminal six histidine and maltose binding protein tandem tags (His×6-MBP). *E. coli* BL21(DE3) cells transformed with this plasmid were grown in LB media, and protein expression was induced as described above. Cells were harvested after overnight and lysed in 20 mM sodium phosphate pH 7.0, 300 mM NaCl. The proteins were purified by Ni-affinity chromatography and SD200 size exclusion chromatography. The His×6-MBP tag were cleaved off by TEV protease and removed by an additional run of SD200 size exclusion chromatography or Source 15S cation chromatography.

### Preparation of the GABARAP∼ATG3_crystal_ conjugate

For large scale preparation of the GABARAP∼ATG3_crystal_ conjugate for crystallization, GABARAP and ATG3_crystal_ were conjugated by mixing 27 µM His×6-MBP-GABARAP, 14 µM ATG3_crystal_ and 1 µM ATG7 in a buffer consisting of 50 mM Hepes pH 7.5, 150 mM NaCl, 1 mM TCEP, 4 mM ATP, 2 mM MgCl_2_. The reaction was allowed to proceed for ∼24 h at 30°C. The His×6-MBP-GABARAP∼ATG3_crystal_ conjugate was purified by Source 15S cation exchange chromatography, Source 15Q anion exchange chromatography, followed by SD200 size exclusion chromatography. The protein was digested with TEV protease and further purified with Source 15Q to remove the His-MBP tag. The GABARAP∼ATG3_crystal_ conjugate was concentrated and the buffer was exchanged to 10 mM HEPES pH 7.0, 150 mM NaCl, and 1 mM DTT.

### Crystallization

A 0.2 µL drop of 10mg/ml GABARAP∼ATG3_crystal_ was mixed with the equal volume of the reservoir solution composed of 0.2 mM ammonium sulfate, 0.1 mM Tris pH8.5, 25%(w/v) PEG3550 and the mixture was placed on a sitting drop plate. The plate was incubated at 4°C. Crystals were equilibrated with a cryoprotection buffer composed of 0.2 mM ammonium sulfate, 0.1 mM Tris pH8.5, 26.5%(w/v) PEG3550, and 20% ethylene glycol, then flash-cooled in liquid nitrogen. Data were collected at the SSRL 11-1 synchrotron beamline equipped with a PILATUS 6M detector.

### **X-** ray data processing and structure determination

The X-ray diffraction data were indexed, integrated and scaled using the XDS software ^67^. The phase problem was solved by molecular replacement using Autosol function of PHENIX package ^68^ and AtATG3 (PDB ID: 3VX8) and GABARAP (PDB ID: 1GNU) crystal structures. The statistics for the data collection and structure determination are listed in Table S1.

### NMR spectroscopy

Isotopically labeled GABARAP and ATG3 proteins were expressed in BL21(DE3) cells grown in M9 minimum media and purified as described above. For the assignment of backbone resonances, 1.2 mM ^15^N/^13^C-labeled ATG3 (Δ89-192) in 20 mM sodium phosphate pH 7.0, 150 mM sodium chloride, 0.5 mM TCEP and 7.5% D_2_O was placed in an NMR tube, and the triple resonance experiments, HNCACB, CBCACONH, HNCO and HNCACO, were recorded on a Bruker 600 MHz spectrometer operated at 40°C. The side-chain resonances of Asn78 were assigned by the absence of the peaks in N78A mutant protein. To monitor the in-trans interaction between free GABARAP and ATG3, 0.15 mM ^15^N-ATG3_nmr_ in a buffer of 20 mM sodium phosphate pH 6.5, 100 mM sodium chloride, 0.5 mM TCEP, and 7.5% D_2_O was placed in an NMR tube, and a mixture of 0.15 mM ^15^N-ATG3_nmr_ and 6 mM GABARAP in the same buffer was titrated in the tube. A ^1^H-^15^N HSQC spectrum was acquired for each titration point at 30°C. The titration-induced changes in chemical shift (Δδ) were calculated using the following equation 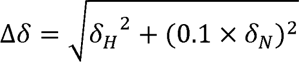. All NMR spectra were processed with NMRPipe and NMRDraw ^69^ and analyzed with NMRViewJ ^70^.

### Conjugation assays

The GABARAP∼ATG3 thioester conjugation and the GABARAP–PE conjugation assays were performed as described previously ^42^. For each thioester conjugation, a reaction mixture containing 1 µM ATG7, 5 µM ATG3, and 7 µM GABARAP was prepared in a buffer of 20 mM HEPES pH7.5, 150 mM NaCl, 1 mM MgCl_2_, and 0.5 mM TCEP. The reaction was initiated by adding 1 mM ATP. At 30, 60, 120 second time points, aliquots were removed from the reaction solution and mixed with 2 x LDS PAGE sample buffer without reducing reagents. At the 120 second, another aliquot was mixed with sample buffer supplemented with 10 % (vol./vol.) β-mercaptoethanol. Samples were electrophoresed at 4°C in homemade 12% acrylamide bis-Tris pH6.5 gels and MOPS buffer pH7.2. Gels were stained with Coomassie Brilliant Blue G-250 and imaged using Odyssey scanner. (Li-Cor), and bands were quantified using the Image Studio Lite software (Li-Cor).

For lipidation assays, liposomes consisting of 40% DOPC (1,2-dioleoyl-sn-glycero-3-phosphocholine), 40% DOPE (1,2-dioleoyl-sn-glycero-3-phosphoethanolamine), 20% bovine liver PI (L-α-phosphatidylinositol) were prepared as previously reported ^42^. In brief, lipids (Avanti Polar Lipids, Inc) were dried under nitrogen gas stream and further vacuumed for 1h. The resulting lipid film was hydrated in 50 mM HEPES pH7.5 and 150 mM NaCl by vigorous mixing for 1h at room temperature and bath-sonicated for 5 min. The hydrated lipids were freeze-thawed three times and extruded through a 50 nm filter membrane. For lipidation, a reaction mixture containing 1 µM ATG7, 1 µM ATG3, 1 µM ATG12-ATG5-ATG16N (ATG16L1 residues 11-43), 10 µM GABARAP and 1 mM liposomes was prepared in a buffer of 20 mM HEPES pH7.5, 150 mM NaCl, 1 mM MgCl_2_, and 0.5 mM TCEP. The reaction was initiated by adding 1 mM ATP, and aliquot were removed from the mixture at each time point and mixed with 2 x SDS-PAGE sample buffer containing 10 % (vol./vol.) β-mercaptoethanol. Samples were electrophoresed at 4°C in 12.5% acrylamide Tris-glycine gels containing 6M urea. Gels were stained and quantitated as described above.

