## Supplemental Materials for "Structural Analyses of a GABARAP∼ATG3 Conjugate Uncover a Novel Non-covalent Ubl-E2 Backside Interaction"

Table S1. Data collection and refinement statistics

| | GBR~ATG3$_{crystal}$ |
|---|---|
| **Data collection** | |
| Wavelength (Å) | 0.9794 |
| Resolution range (Å) | 38.33–2.7 (2.796–2.7) |
| Space group | $P4_32_12$ |
| Unit cell | |
| *a*, *b*, *c* (Å) | 97.388, 97.388, 166.774 |
| α, β, γ (°) | 90, 90, 90 |
| Total reflections | 229322 (21522) |
| Unique reflections | 22689 (2200) |
| Redundancy | 10.1 (9.8) |
| Completeness (%) | 99.56 (99.46) |
| $R_{merge}$ | 0.06022 (1.232) |
| $I / \sigma I$ | 24.97 (1.95) |
| $R_{merge}$ | 0.06022 (1.232) |
| CC1/2 | 1 (0.764) |
| CC* | 1 (0.931) |
| **Refinement** | |
| Initial models | PDB IDs: 3VX8, 1GNU |
| Reflections used in refinement | 22652 (2200) |
| Reflections used for R-free | 1994 (193) |
| $R_{work}$ | 0.2191 (0.3662) |
| $R_{free}$ | 0.2481 (0.3880) |
| Number of non-hydrogen atoms | |
| Protein | 4846 |
| Ligand/ion | 21 |
| Water | 10 |
| Protein residues | 582 |
| *B*-factors | |
| Protein | 89.49 |
| Ligand/ion | 86.78 |
| Water | 61.70 |
| R.m.s. deviations | |
| Bond lengths (Å) | 0.002 |
| Bond angles (°) | 0.48 |
| Ramachandran plot | |
| Favored (%) | 96.32 |
| Allowed (%) | 3.68 |
| Outliers (%) | 0.00 |
| Rotamer outliers (%) | 0.95 |
| Clashscore | 3.92 |
| MolProbity score | 1.43 |

Values in parentheses are for the highest-resolution shell. Each data set was collected from one crystal.

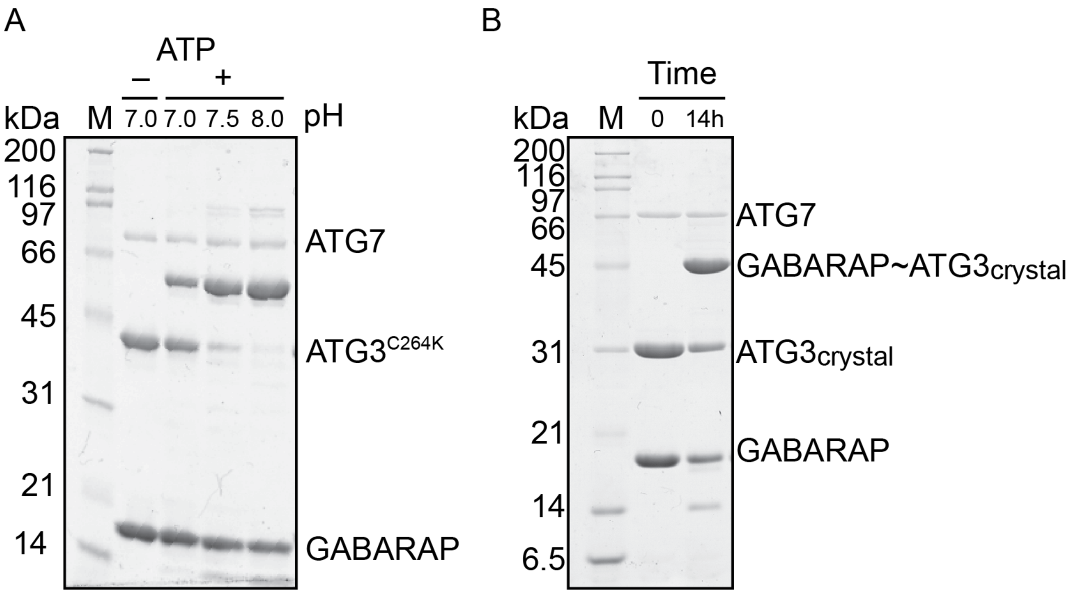


**Figure S1.** SDS-PAGE analysis of the GABARAP~ATG3$^{C264K}$ conjugation. (A) Conjugation reactions using full-length ATG3$^{C264K}$. The reaction mixtures containing 30 μM GABARAP, 15 μM ATG3$^{C264K}$, 1 μM ATG7, 5 mM ATP, 2 mM $MgCl_2$, 50 mM HEPES (pH7.0, 7.5, or 8.0), 150 mM NaCl, and 1 mM TCEP were incubated for 24 h at 30˚C. The control reaction was performed without ATP at pH7.0. SDS-PAGE samples were prepared by mixing the reaction solutions with 2 x SDS-PAGE sample buffer containing 10 % (vol./vol.) β-mercaptoethanol. (B) A conjugation reaction with ATG3$_{crystal}$. The reaction was performed under the same condition as in (A) except that the buffer pH was 7.5, and the incubation time was 14 h.

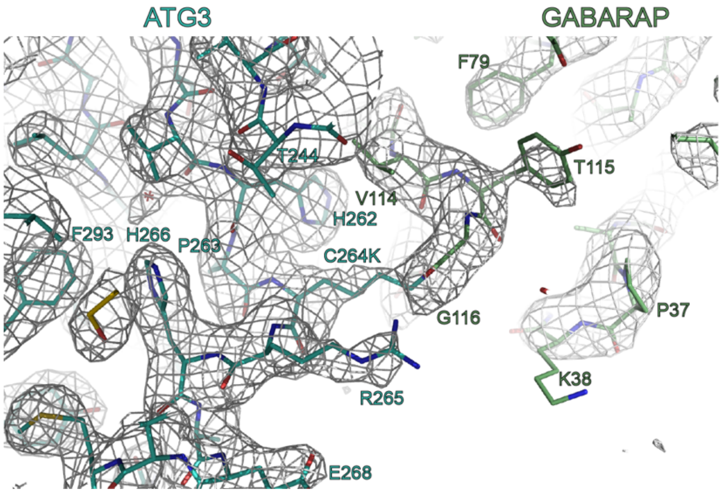


**Figure S2.** The $2mF_0 - DF_c$ electron density map around the isopeptide conjugate site between GABARAP and ATG3. The map is contoured at 1σ.

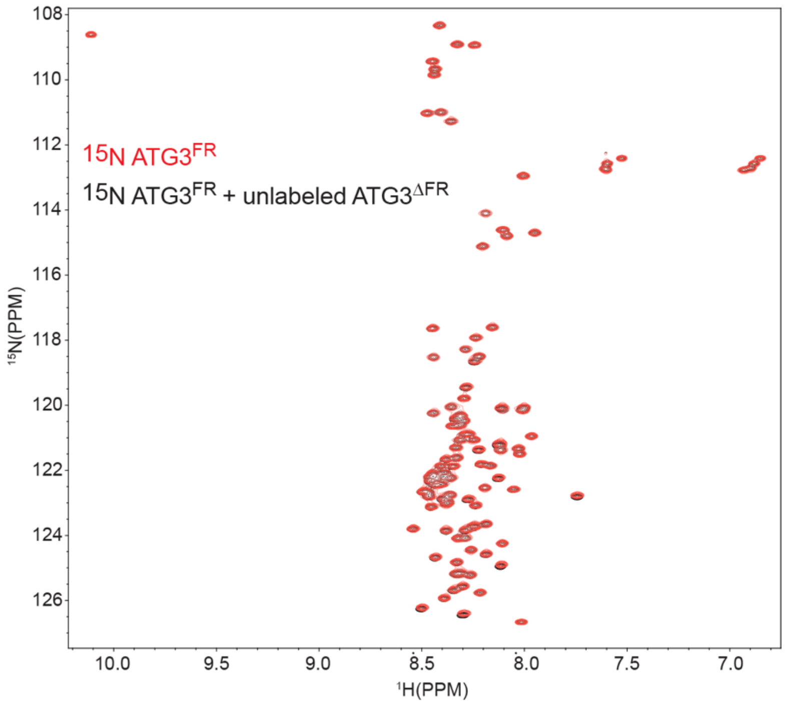


**Figure S3.** Overlay of $^{1}$H-$^{15}$N HSQC spectra of 140 μM $^{15}$N-labeled ATG3$^{FR}$ (residues 92–192) with (red) and without (black) 350 μM unlabeled ATG3$^{\Delta FR}$ (residues 1–91 fused to 193-314).

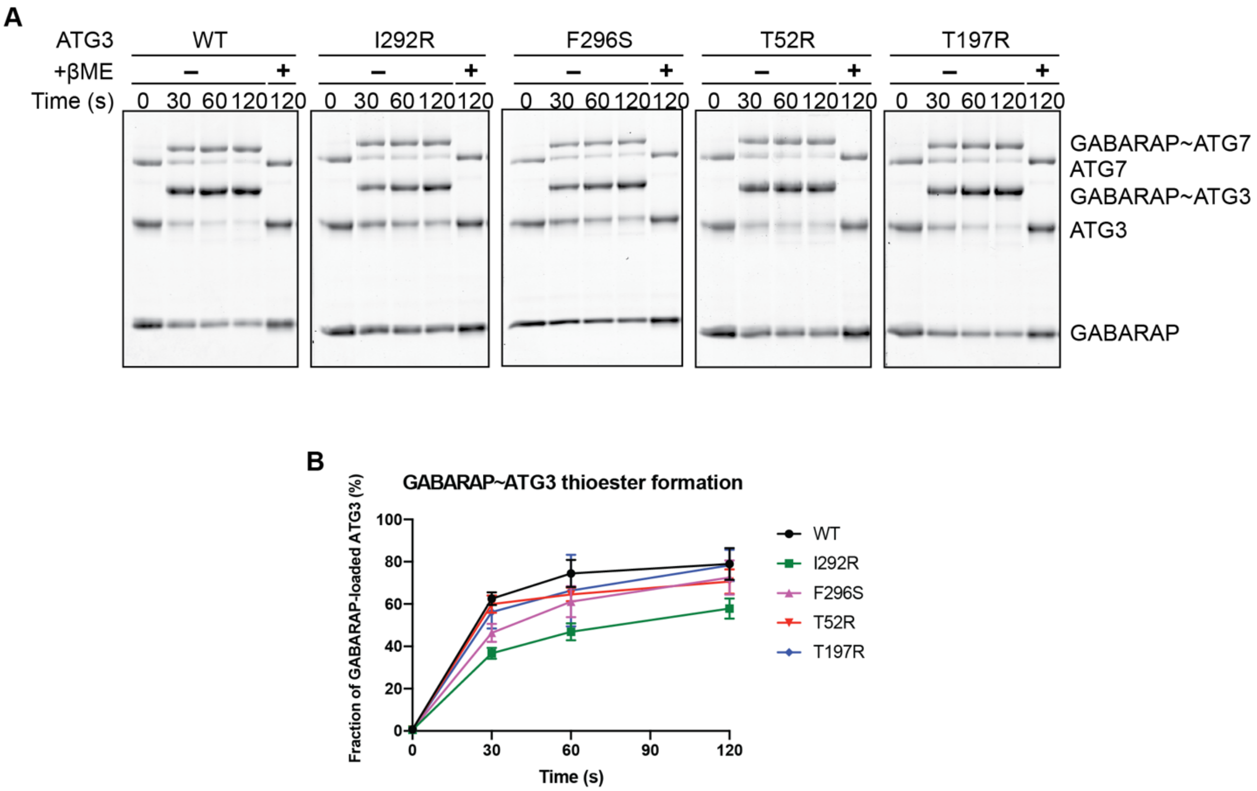


**Figure S4.** GABARAP loading of ATG3 mutants. (A) Neutral pH SDS-PAGE analyses of the GABARAP~ATG3 thioester formation with ATG3 mutants. (B) Quantification of (A). The error represents S.D. (n=3).

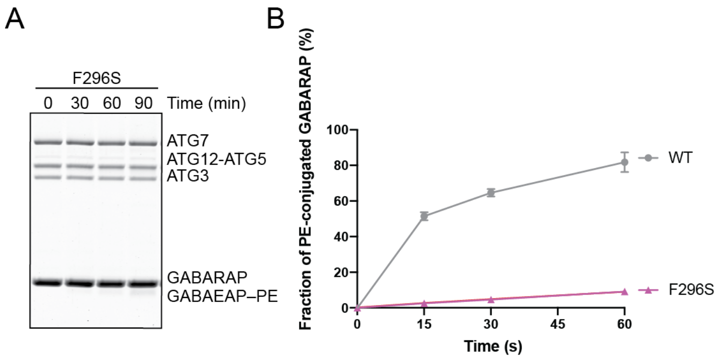


**Figure S5.** GABARAP–PE conjugation assay with the ATG3 F296S mutant. (A) SDS-PAGE analysis of the reaction. (B) Quantification of the reaction. Data are shown in the same manner as in Fig. 1E. The WT data are the same as in Fig. 1E.

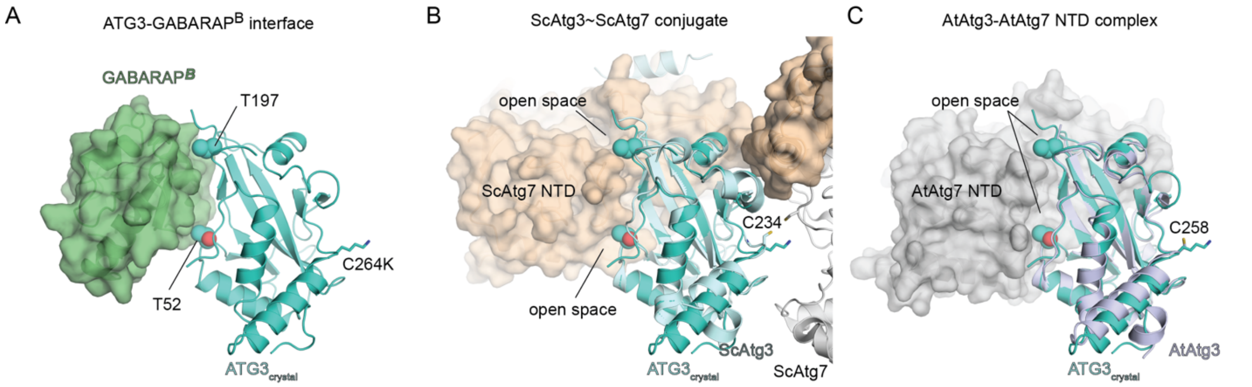


**Figure S6.** Positions of Thr52 and Thr197 of ATG3 with respect to $GABARAP^B$ and the E1 NTDs. (A) The $ATG3_{crystal}$-$GABARAP^B$ unit in the crystal structure. There is little open space around the two threonine residues within the interface between the two proteins. (B) Superimposition of $ATG3_{crystal}$ onto the ScAtg3~ScAtg7 disulfide-crosslinked conjugate (PDB ID: 4GSL). Both T52 and T197 of ATG3 would have open space toward the outside of the ATG7 NTD. (C) Superimposition of $ATG3_{crystal}$ onto the AtAtg3-AtAtg7 NTD complex (PDB ID: 3VX8). T197 would face toward the outside of the ATG7 NTD, while T52 would have much open space in the cavity of the ATG7 NTD interface.
